# A convolutional neural network highlights mutations relevant to antimicrobial resistance in *Mycobacterium tuberculosis*

**DOI:** 10.1101/2021.12.06.471431

**Authors:** Anna G. Green, Chang H. Yoon, Michael L. Chen, Luca Freschi, Matthias I. Gröschel, Isaac Kohane, Andrew Beam, Maha Farhat

## Abstract

Long diagnostic wait times hinder international efforts to address multi-drug resistance in *M. tuberculosis*. Pathogen whole genome sequencing, coupled with statistical and machine learning models, offers a promising solution. However, generalizability and clinical adoption have been limited in part by a lack of interpretability and verifiability, especially in deep learning methods. Here, we present a deep convolutional neural network (CNN) that predicts the antibiotic resistance phenotypes of *M. tuberculosis* isolates. The CNN performs with state-of-the-art levels of predictive accuracy. Evaluation of salient sequence features permits biologically meaningful interpretation and validation of the CNN’s predictions, with promising repercussions for functional variant discovery, clinical applicability, and translation to phenotype prediction in other organisms.

## 1. Introduction

Tuberculosis is a leading cause of death worldwide from an infectious pathogen, with more than 1.5 million people succumbing to the disease annually(*1*). Rising rates of antibiotic-resistant *Mycobacterium tuberculosis*, the causative agent of tuberculosis, continue to rise, pose a threat to public health(*2*). A major challenge in combatting antibiotic-resistant tuberculosis is the timely selection of appropriate treatments for each patient, particularly when growth-based drug susceptibility testing takes weeks(*1*).

Molecular diagnostic tests for *M. tuberculosis* antimicrobial resistance reduce the time to result to hours or days, but only target a small number of loci relevant to a few antibiotics, and cannot detect most rare genetic variants(*3*). Although whole genome sequencing-related diagnostic tests offer the promise of resolving some of these deficiencies, statistical association techniques have seen limited success, hindered by their inability to assess newly observed variants and epistatic effects(*3*–*7*). More complex models such as deep learning provide promising flexibility but are often uninterpretable, making them difficult to audit for safety purposes (*8*, *9*). Moreover, interrogating black box models offers the opportunity for hypothesis generation which can be later validated, potentially improving scientific understanding of the underlying phenomenon (*10*).

A recent “wide-and-deep” neural network applied to *M. tuberculosis* genomic data outperformed previous methods to predict antimicrobial resistance to 10 antibiotics(*11*); however, like most deep learning methods, the logic behind its predictions was indiscernible. Although more interpretable rule-based classifiers of antimicrobial resistance in *M. tuberculosis* have been developed(*12*, *13*), these rely on predetermined single-nucleotide polymorphisms or *k*-mers, hindering their flexibility to generalize to newly observed mutations, and universally ignore genomic context. Deep convolutional neural networks (CNNs), which greatly reduce the number of required parameters compared to traditional neural networks, could be used to consider multiple complete genomic loci with the ultimate goal of incorporating the whole genome. This would allow the model to assess mutations in their genetic context by capturing the order and distance between resistance mutations of the same locus, allowing a better incorporation of rare or newly observed variation. Deep CNNs, when paired with attribution methods that highlight the most salient features informing the model predictions, are a promising means of harnessing the predictive power of deep neural networks in genomics for biological discovery and interpretation(*14*). CNNs also have the added advantage of minimizing the preprocessing needed of genomic variant data. The extent to which we may trust these highlighted features remains the subject of ongoing scientific exploration(*8*, *15*, *16*).

Here, we show that CNNs perform *en par* with the state-of-the-art in predicting antimicrobial resistance in *M. tuberculosis* and provide biological interpretability through motif representation captured in saliency mapping. We train two models: one designed for accuracy that incorporates genetic and phenotypic information about all drugs; and a second designed for interpretability that forces the model to only consider putatively causal regions for a particular drug. Our models are trained on the entire genetic sequence of 18 regions of the genome known or predicted to influence antibiotic resistance, using data collected from over 20,000 *M. tuberculosis* strains spanning the four major global lineages. Across each locus, we calculate genomic positions that most influence the prediction of resistance for each drug, validating our method by recapitulating known positions and providing predictions of new positions potentially involved in drug resistance. Given the growing movement towards greater interpretability in machine learning methods(*16*, *17*), we expect this model to have implications for hypothesis generation about molecular mechanisms of antimicrobial resistance through genotype-phenotype association.

## 2. Results

### Training dataset characteristics

We train and cross-validate our models using 10,201 *M. tuberculosis* isolates from the ReSeqTB and the WHO Supranational Reference Laboratory Network (sources detailed in the **Materials and Methods)**. Each isolate is phenotyped for resistance to at least one of thirteen antitubercular drugs: the four first-line drugs isoniazid, rifampicin, ethambutol, and pyrazinamide, and nine additional second-line drugs (**Table 1**). All drugs are represented by at least 250 phenotyped isolates.

**Table 1a:**
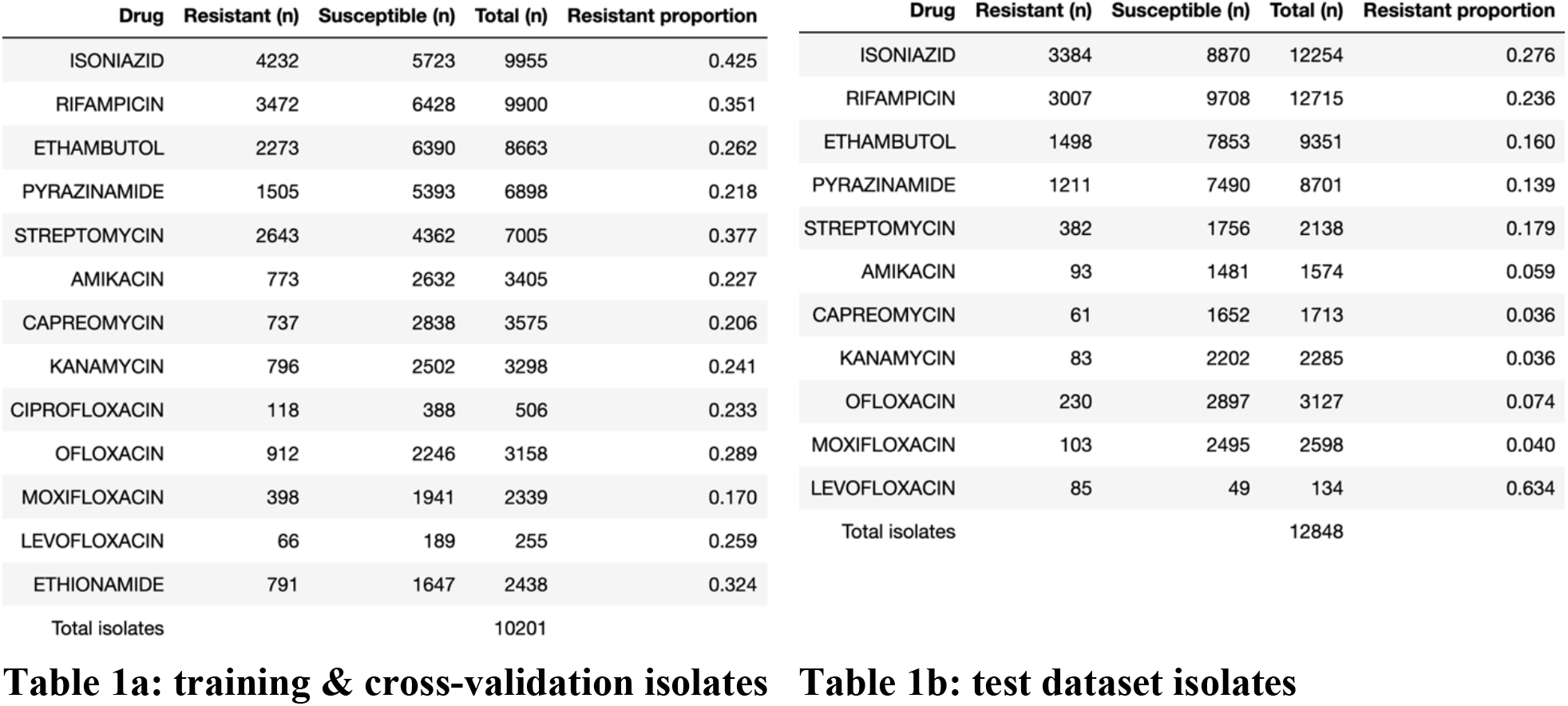
**Tables 1a & 1b**: Phenotypic summary of the 23,049 isolates used to train and cross-validate (1a), and test (1b) the models: the numbers of resistant isolates, susceptible isolates, the total tested (sum of the numbers of resistant and susceptible isolates), and the resistant proportion, with respect to each of the 13 anti-TB drugs (training and cross-validation) or 11 anti-TB drugs (test). Ciprofloxacin and ethionamide were excluded from the test dataset due to small numbers (0/2 resistant to ciprofloxacin; 12/25 resistant to ethionamide).

### Model design

We build two models to predict antibiotic resistance phenotypes from genome sequences. The first is a multi-drug convolutional neural network (MD-CNN), designed to predict resistance phenotypes to all 13 drugs at once. The model inputs are the full sequences of 18 loci in the *M. tuberculosis* genome, selected based on known or putative roles in antibiotic resistance **(Table 2)**. We chose the final MD-CNN architecture using an iterative process **(Figure 1, Supplementary Figure 1)**. As superior performance of multi-task over single-task models has been demonstrated with convolutional neural networks in computer vision(*18*–*20*), the MD-CNN is designed to optimize performance by combining all genetic information and relating it to the full resistance antibiogram. We compare the MD-CNN with 13 single-drug convolutional neural networks (SD-CNN), each of which has a single-task, single-label architecture, in which only loci with previously known causal associations for any given drug are incorporated **(Supplementary Figure 2)**. We benchmark both types of CNNs against an existing state-of-the-art multi-drug wide-and-deep neural network (MD-WDNN)(*11*), and a logistic regression with L2 regularization penalty.

**Table 2:** Loci included in the MD-CNN and SD-CNN models. The 18 loci included in the MD-CNN and their start and end coordinates (in H37Rv numbering). Each locus was designated as putatively involved in resistance to at least one drug. To construct the 13 SD-CNN models, the relevant loci for each drug were combined – for example, the isoniazid (INH) model contained the *acpM-kasA, oxyR-ahpC, katG, and fabG1-inhA* loci.

| **Locus** | **Start** | **End** | **Drug(s)** | **Length (in H37Rv)** |
| --- | --- | --- | --- | --- |
| *acpM-kasA* | 2517695 | 2519365 | Isoniazid | 1670 |
| *gid* | 4407528 | 4408334 | Streptomycin | 806 |
| *rpsA* | 1833378 | 1834987 | Pyrazinamide | 1609 |
| *clpC* | 4036731 | 4040937 | Pyrazinamide | 4206 |
| *embCAB* | 4239663 | 4249810 | Ethambutol | 10147 |
| *aftB-ubiA* | 4266953 | 4269833 | Ethambutol | 2880 |
| *rrs-rrl* | 1471576 | 1477013 | Streptomycin, Amikacin, Capreomycin, Kanamycin | 5437 |
| *ethAR* | 4326004 | 4328199 | Ethionamide | 2195 |
| *oxyR-ahpC* | 2725477 | 2726780 | Isoniazid | 1303 |
| *tlyA* | 1917755 | 1918746 | Capreomycin | 991 |
| *katG* | 2153235 | 2156706 | Isoniazid | 3471 |
| *rpsL* | 781311 | 781934 | Streptomycin | 623 |
| *rpoBC* | 759609 | 767320 | Rifampicin | 7711 |
| *fabG1-inhA* | 1672457 | 1675011 | Isoniazid, Ethionamide | 2554 |
| *eis* | 2713783 | 2716314 | Kanamycin, Amikacin | 2531 |
| *gyrBA* | 4997 | 9818 | Ciprofloxacin, Levofloxacin, Moxifloxacin, Ofloxacin | 4821 |
| *panD* | 4043041 | 4045210 | Pyrazinamide | 2169 |
| *pncA* | 2287883 | 2289599 | Pyrazinamide | 1716 |

**Figure 1:**
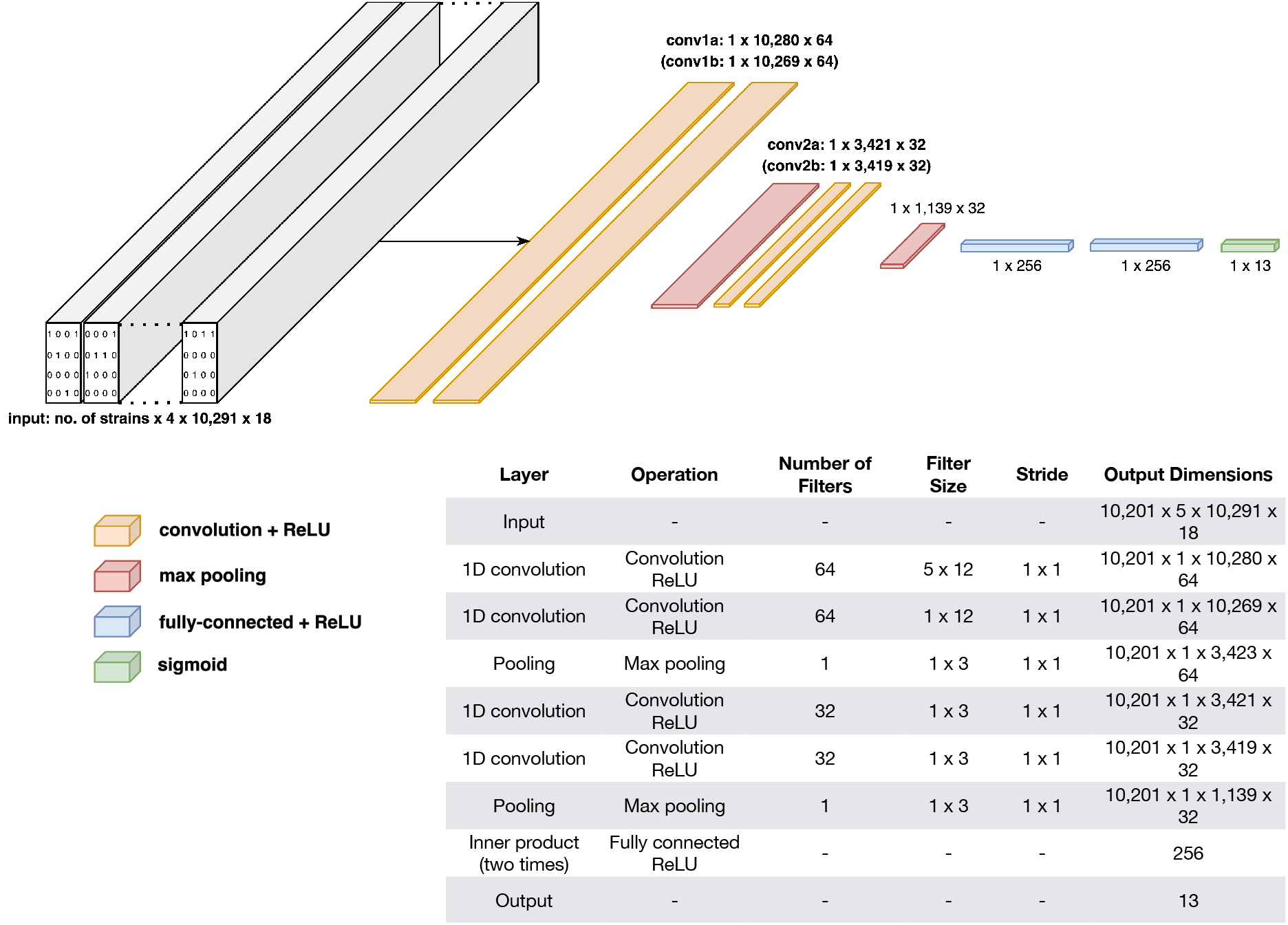
schematic diagram and table of the multi-drug convolutional neural network (MD-CNN). In the output layer, each of the 13 nodes is composed of a sigmoid function to compute a probability of resistance for their respective anti-TB drug (13 anti-TB drugs in total). The input consisted of ‘10,201’ isolates (TB strains) for which there was resistance phenotype data for at least 2 anti-TB drugs; ‘5’ for one-hot encoding of each nucleotide (5 dimensions, one for each nucleotide – adenine, thymine, guanine, cytosine plus gaps); ‘10,291’ being the number of nucleotides of the longest locus (*embC-embA-embB*); ‘18’ loci of interest were incorporated as detailed in ‘Materials and methods’.

### Benchmarking CNN models against state-of-the-art

We used 5-fold cross-validation to compare the performance of the four architectures (MD-CNN, the SD-CNN, L2 regression, and WDNN(*11*)) on the training dataset (N=10,201 isolates, **Supplementary Table 1**).

The mean MD-CNN AUC of 0.912 for second-line drugs is significantly higher than the mean 0.860 for L2 regression (Welch’s t-test with Benjamini-Hochberg FDR *q*<0.05), but the mean AUCs for first-line drugs (0.948 *vs*. 0.923) are not significantly different (Benjamini-Hochberg *q*=0.055). The mean SD-CNN AUCs of 0.938 (first-line drugs) and 0.877 (second-line drugs) are not significantly different than for L2 regression (first-line *q*=0.20, second-line *q* = 0.16). However, L2 regression demonstrates much wider confidence intervals than the CNN models (median 0.037 versus 0.010, IQR 0.035 versus 0.014), indicating a lack of reliability as the performance depends on the particulars of the cross-validation split (**Figure 2)**.

**Figure 2:**
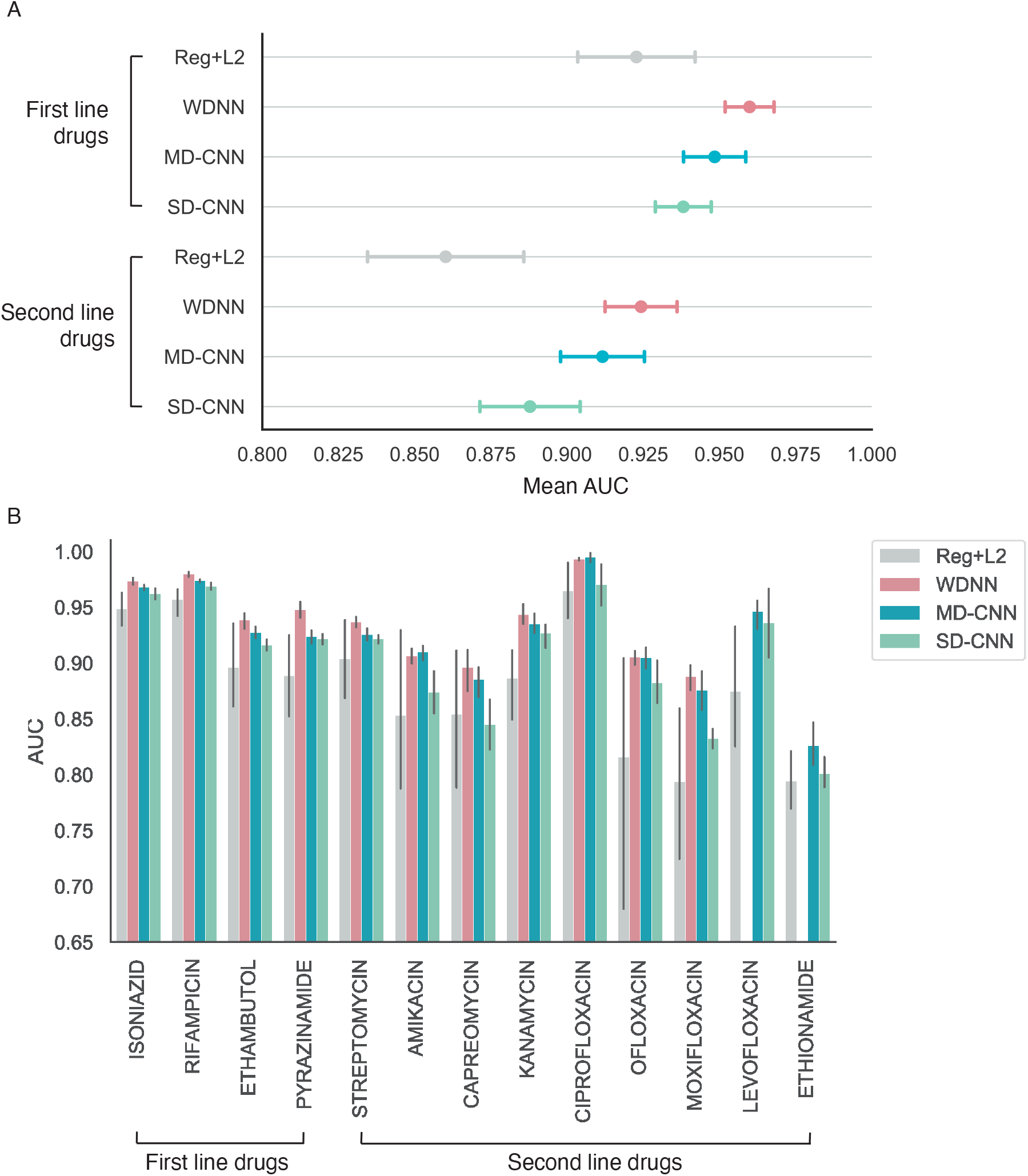
**MD-CNN performs comparably to state-of-art WDNN** for both first- and second-line drugs. Results of five-fold cross validation on the training dataset for the four models: WDNN, logistic regression + L2 benchmark, SD-CNN, and MD-CNN. (A) mean AUC and 95% confidence intervals, pooled for first and second line drugs. (B) mean AUC and 95% confidence intervals for each drug. The WDNN was not initially trained on levofloxacin or ethionamide and thus was not evaluated for these drugs.

Against the state-of-the-art WDNN, the AUCs, sensitivities, and specificities of the MD-CNN are comparable: the MD-CNN’s mean AUC is 0.948 (vs. 0.959 for the MD-WDNN, *q*=0.15) for first-line drugs, and 0.912 (vs. 0.924 for the MD-WDNN, *q*=0.30) for second-line drugs. The SD-CNN is less accurate than the MD-WDNN for both first-line (Benjamini-Hochberg *q*=0.006) and second-line drugs (q = 0.005, **Supplementary Table 1, Figure 2)**.

The SD-CNN (mean AUC of 0.938 for first-line drugs; mean AUC of 0.877 for second-line drugs) performs comparably to the MD-CNN for first-line drugs (*q*=0.19), and is less accurate than the MD-CNN for second-line drugs (*q*=0.009).

### CNN models generalize well on hold-out test data

We test the generalizability and real-world applicability of our CNN models on a holdout dataset of 12,848 isolates which were curated on a rolling basis during our study (**Table 1b, Materials and Methods**). Rolling curation provides a more realistic test of generalizability to newly produced datasets. Due to rolling curation and source difference, the test dataset exhibits different proportions of resistance to the 13 drugs (e.g. isoniazid resistance in 28% vs. 43% in the training dataset). We assessed generalizability of the models using phenotype data for 11 drugs in the hold-out test dataset, since it contained low resistance counts for ciprofloxacin and ethionamide.

We find that the MD-CNN generalizes well to never-before-seen data for first-line antibiotic resistance prediction, achieving mean AUCs of 0.965 (95% confidence interval [C.I.] 0.948 - 0.982) on both training and hold-out test sets for first-line drugs **(Figure 3)**. However, generalization for second-line drugs is mixed: for the drugs streptomycin, amikacin, ofloxacin, and moxifloxacin, the model generalizes well, achieving mean AUCs of 0.939 (CI 0.928 - 0.949) on the test data (compared with 0.939 (CI 0.929 - 0.949) on the training data). For the second-line drugs capreomycin, kanamycin, and levofloxacin, the model generalization was reduced, achieving mean AUCs of 0.831 (CI 0.824 - 0.838) on the test data (compared with 0.955 (CI 0.931 – 0.978) on the training data). We find that the SD-CNN generalizes well on first-line drug resistance for hold-out test data, with a mean AUC of 0.956 (CI 0.929 – 0.974). The SD-CNN also generalizes well for second-line drugs, with a mean AUC of 0.862 (CI 0.830 – 0.894).

**Figure 3:**
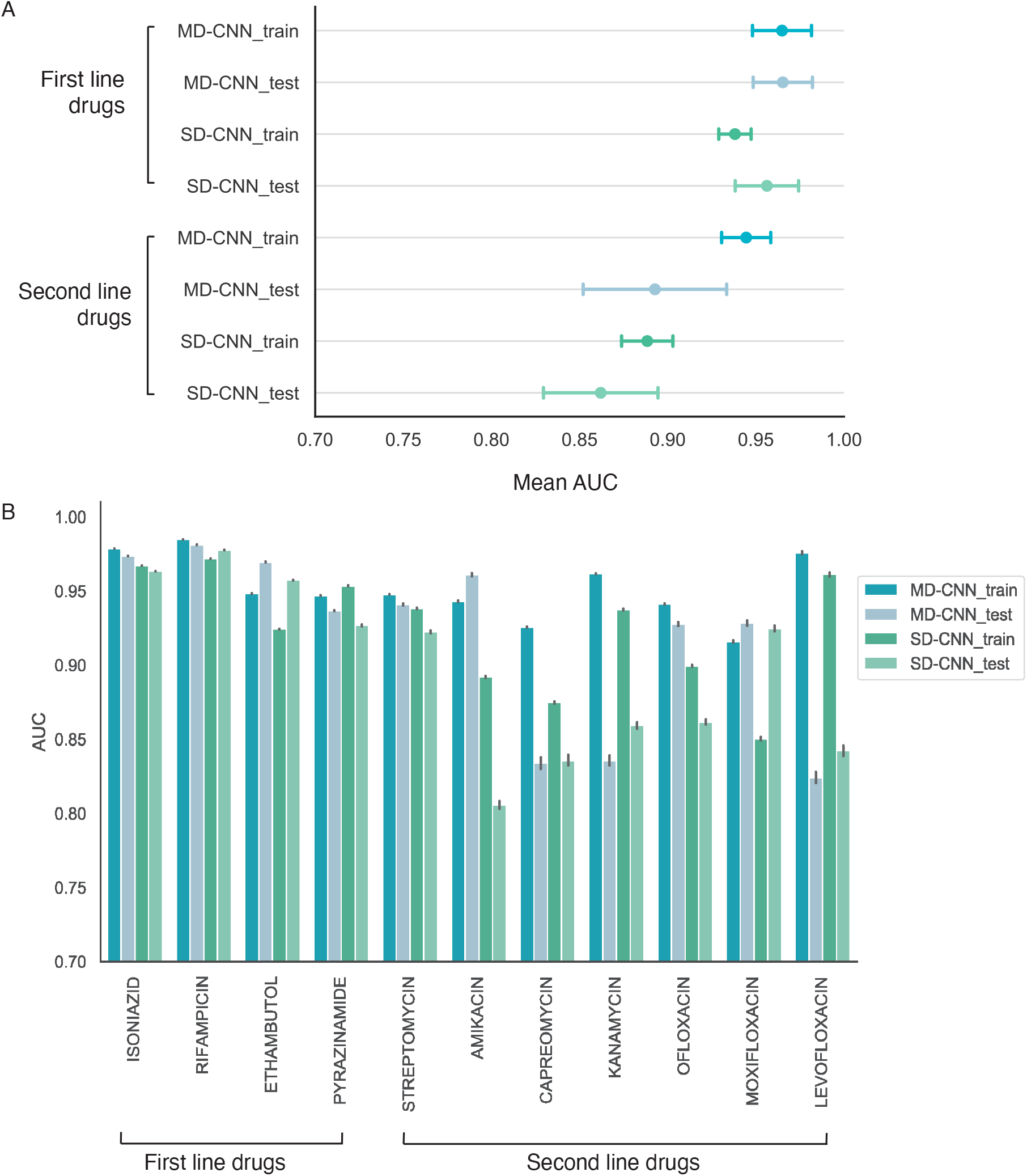
MD-CNN and SD-CNN model generalize well on hold-out test data. Performance of CNN models trained on the entire training dataset evaluated on either the training dataset or the hold-out test dataset. (A) Mean AUC and 95% confidence intervals (calculated across drugs) for first- and second-line drugs, pooled. (B) Mean AUC for each drug with confidence intervals generated by 100x bootstrapping with 80% of isolates. Ciprofloxacin and ethionamide were not assessed due to low number of resistant isolates.

We test the hypothesis that missed resistance (false negatives) is due to mutations affecting phenotype found outside of the 18 incorporated loci. To achieve this, we compute the number of mutations in the incorporated loci that separate each test isolate from the nearest isolate(s) in the training set and the corresponding phenotype of the nearest isolates **(Methods)**. We find that many of the false negatives have a genomically identical yet sensitive isolate in the training set, ranging from a minimum of 34% for pyrazinamide to a maximum of 86% for kanamycin, and suggesting that additional mutations outside of the examined loci may influence the resistance phenotype.

### MD-CNN achieves accuracy by learning dependency structure of drug resistances

Because the inputs to the CNN models are the complete sequence of 18 genetic loci involved in drug resistance, we are able to assess the contribution of every site, in its neighboring genetic context, to the prediction of antibiotic resistance phenotype. We do this by calculating an importance score for each nucleotide site in each input sequence using DeepLIFT(*21*). For any input, DeepLIFT calculates the change in predicted resistance relative to a reference input, and then backpropagates that difference through all neurons in the network to attribute the change in output to changes in the input variable. We use the pan-susceptible H37Rv genome as a reference(*22*). We take the highest magnitude (positive or negative) importance score for each nucleotide across all isolates in the training set (**Methods**).

We find evidence that the MD-CNN achieves high performance by relying on drugdrug resistance correlations. Due to the global standard therapeutic regimen for tuberculosis, resistances to first-line drugs almost always evolve before resistances to second-line drugs, and frequently in a particular order(*23*) **(Figure 4A-B)**. When considering the top 0.01% (N=17) of positions with the highest DeepLIFT importance scores for each drug, we observe that an average of 85.0% are known to confer resistance to any drug(*24*), but only a mean of 24.0% are known to confer resistance to the particular drug being investigated. For example, the top three hits for the antibiotic kanamycin are, in order, a causal hit to the *rrs* gene, an ethambutol-resistance causing hit to the *embB* gene, and a fluoroquinolone-resistance-causing hit to the *gyrA* gene (**Extended Data 1)**. To probe this further, we introduce mutations that confer resistance to the first-line drugs rifampicin and isoniazid into a pan-susceptible genomic sequence background, *in silico*, and this increased the MD-CNN predicted resistance probability of pyrazinamide, streptomycin, amikacin, moxifloxacin and ofloxacin resistance (**Figure 5A**). The MD-CNN model generalized well for all five of these drugs: AUC of 0.939 for these drugs versus 0.831 for the remaining second-line drugs. Taken together, these observations show that the MD-CNN benefits from the correlation structure of antibiotic resistance.

**Figure 4:**
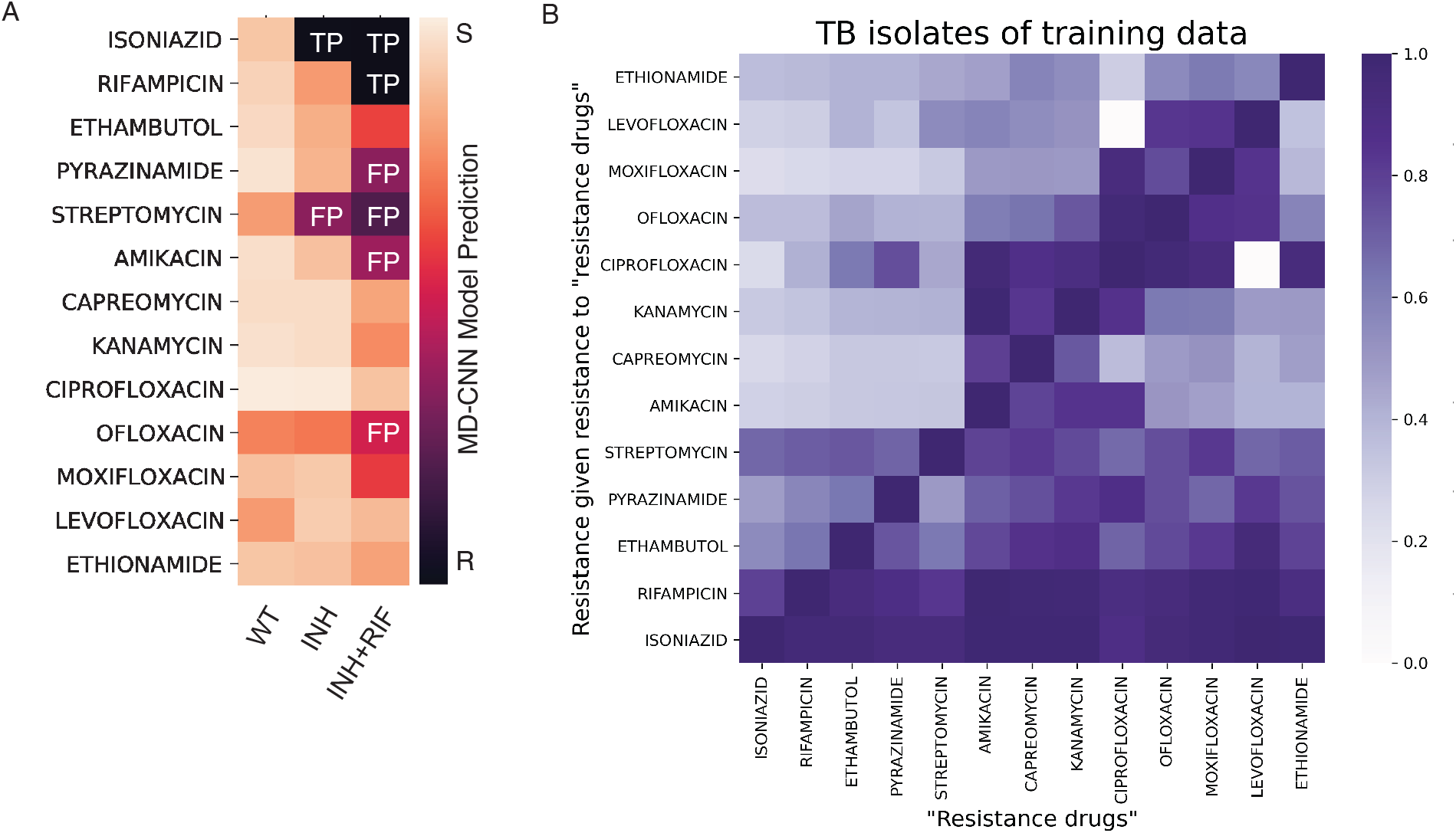
MD-CNN learns dependency structure of antibiotic resistance. **(A)** Introduction of single resistance-conferring mutations into pan-susceptible wild-type background (H37Rv) is sufficient to cause MD-CNN model to predict false positive resistances. A single isoniazid-resistance conferring mutations (2155168G, *katG* S315T) or one isoniazid- and one rifampicin-resistance conferring mutation (2155168G and 761155T, *rpoB* S450L) were introduced *in silico* into the wild-type background sequence and resistances were predicted using the MD-CNN model. (B) Dependency heatmaps of drug resistance for training isolates. The horizontal axis represents the drugs to which isolates exhibited resistance. Based on this condition of resistance, the proportion of resistance to other drugs (vertical axis) was computed.

**Figure 5:**
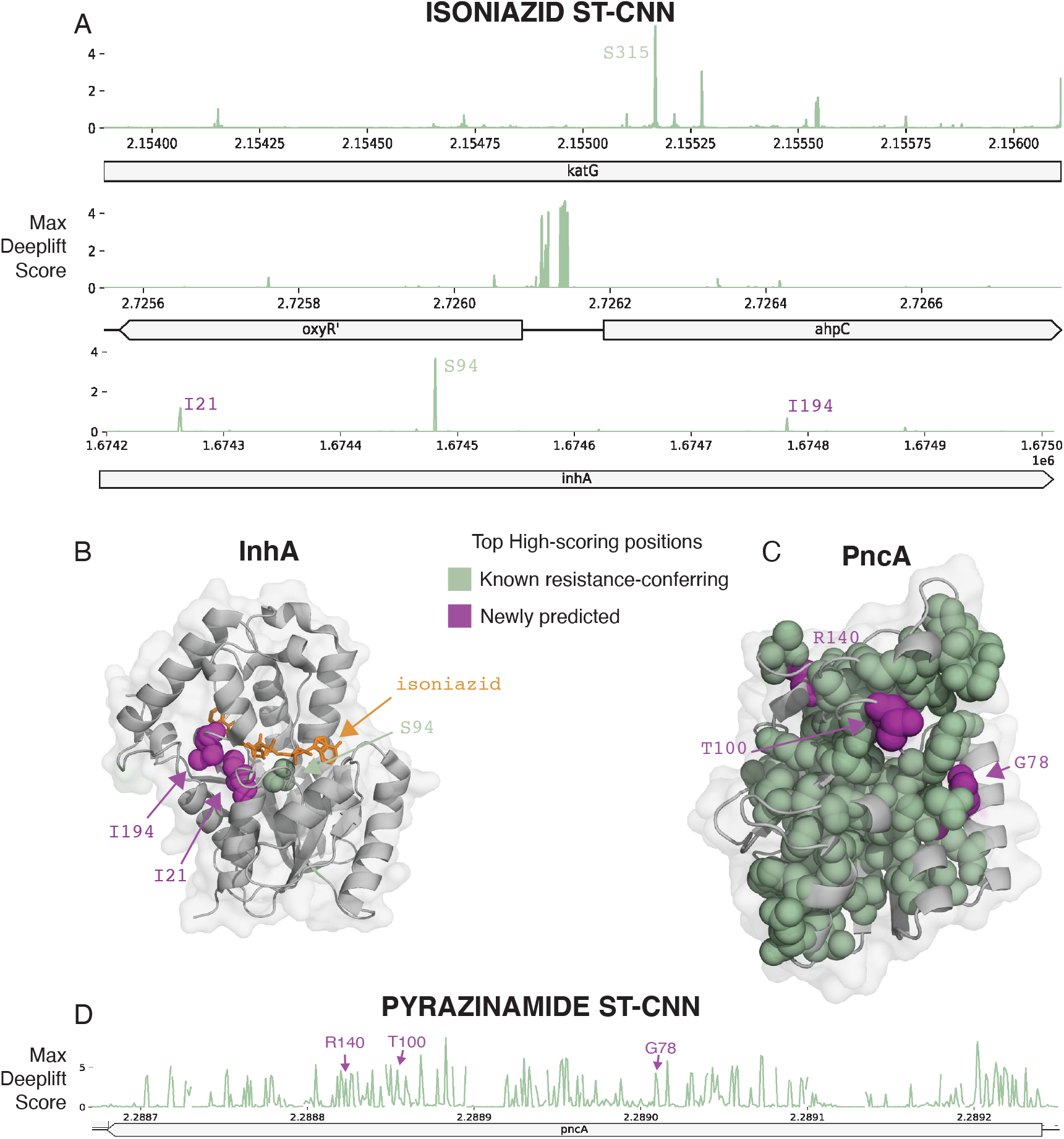
SD-CNN importance scores highlight known and plausible new resistance-conferring loci. Variants not known to cause resistance according to the WHO(*24*) are shown in purple. (A) Maximum of absolute value DeepLIFT importance scores for the isoniazid SD-CNN across all isoniazid-resistant loci. (B) High-importance variants in the InhA protein mapped to its crystal structure(*60*). (C) High-importance variants in the PncA protein mapped to its crystal structure(*61*). (D) Maximum of absolute value DeepLIFT importance scores for the pyrazinamide SD-CNN in the *pncA* locus.

### SD-CNN saliencies highlight known and new potential predictors of resistance

We assess whether the DeepLIFT saliency scores for the SD-CNN models are able to capture known causal, resistance-conferring variants by cross-referencing the WHO catalog of established resistance-conferring mutations(*24*). We find that of the 0.1% of sites with the largest absolute DeepLIFT saliencies in each model, a large proportion are in the WHO catalog of known resistance-conferring positions (ranging from 37.5% for streptomycin to 100% for capreomycin, **Methods, Supplementary Table 2**). In total, we identify 38 variants in the top 0.1% of sites that are not previously known to cause resistance, or classified by the WHO as of “uncertain significance”. Variants associated with the *M. tuberculosis* population structure comprise a smaller proportion, ranging from 0% to 8% of the top 0.1% of hits for each locus (**Methods, Supplementary Table 3**). We examine the distribution of saliency scores closely for two drugs with well understood resistance mechanisms: rifampicin and isoniazid; and for pyrazinamide a drug for which elucidating resistance mechanisms has been more challenging.

#### Rifampicin

Positions in the *rpoB* gene known to cause rifampicin resistance(*24*) constitute 86% of the top 0.1% and 55% of the top 1% of saliency scores **(Supplementary Figure 4)**. Four of the five highest-scoring variants that have not been previously identified as resistance-causing are located in three-dimensional proximity (minimum atom distance < 8Å) to resistance-conferring variants in the RpoB protein structure, demonstrating the biological plausibility for these newly identified sites to confer resistance **(Supplementary Figure 4)**.

#### Isoniazid

The common causal site KatG S315 has the highest maximum saliency in the isoniazid SD-CNN **(Figure 5A)**. We observe several high saliency peaks in the promoter region of the *ahpC* gene, which are currently designated as “uncertain significance” to isoniazid resistance by the WHO(*25*). We observe three saliency peaks in the InhA protein, the mycolic acid biosynthesis enzyme targeted by isoniazid. One peak was at the known resistance-conferring mutation S94, and two at positions I21 and I194, of uncertain significance in the WHO catalogue. All three of these positions are close in 3D structure (minimum atom distance <8Å) to the bound isoniazid molecule(*26*). **(Figure 6B)**

#### Pyrazinamide

Of the top 1% of high saliency positions, 62% are known to be resistance-conferring, and an additional 23% are in *pncA*, but not previously known to cause resistance. The top three of these unknown *pncA* mutations are physically adjacent to known resistance-conferring mutations (**Figure 5 C,D)**. The top 1% of salient positions also includes positions in *clpC1*, a gene recently implicated in pyrazinamide resistance, but mutations within are not yet recognized to be useful for resistance prediction(*27*, *28*) **(Extended Data 1)**.

## 3. Discussion

In summary, we find that the convolutional neural networks offer similar predictive accuracy to the state-of-art MD-WDNN while also being able to discover new loci implicated in resistance, and to visualize them in their genomic context. Another major advantage of the CNNs, is that they require significantly less pre-processing and curation because they directly analyze alignments of genomic loci, allowing the models to consider not only single nucleotide polymorphisms but also sequence features such as insertions and deletions or more complex variation. They also circumvent challenges arising from differing variant naming conventions, and in reconciling variant features across datasets and time.

We find the MD-CNN’s AUCs to be similar to those of the drug-specific SD-CNNs for first-line drugs, and are significantly higher for second-line drugs. CNNs generalize well to the distinct, hold-out, test isolates for these first-line antibiotics, a promising aspect if they are to be deployed in clinical practice. By contrast, there are more mixed results and generally lower hold-out test AUCs for second-line drugs. For both first- and second-line drugs, we observe that false negative isolates are often genetically identical at the considered loci to their drug-sensitive counterparts in the training dataset, indicating that additional genetic information is needed to accurately predict the phenotype for certain isolates.

Although deep neural networks are generally deemed to be less interpretable than traditional statistical methods, we are able to apply two distinct methods to interpret the network’s inner logic: first, assessing model predictions using *in silico* mutagenesis; and second, assessing DeepLIFT importance scores for every input site. By computationally introducing resistance-conferring mutations into known susceptible sequences, we discover that the MD-CNN’s predictions for second-line drugs relies on the correlation structure of drug resistance which is present in both the training and test set. Using DeepLIFT, we highlight which sequence features are informing model predictions: for example, our model confirmed the importance of known, resistance-conferring mutations, such as in the *rpoB*, and *katG* genes.

In addition to highlighting known resistance-conferring mutations, our model discovers 38 resistance variants previously unknown or of “uncertain significance” based on the WHO catalogue(*24*). Including these mutations in resistance prediction may be useful for clinical diagnosis of antibiotic resistance – for example, 6% of isoniazid resistant strains contain at least one newly discovered mutation, and 2.4% contain only newly discovered mutations and no canonical resistance variants. The interpretable, nucleotide-level saliency scores permit the protein contextualization of mutations and offers the prospect of modeling how certain mutations would impact protein structure, and drug binding. This can allow for *in silico* prioritization of putative mutations for further experimental validation.

Limitations of this study include: first, the genomic variants highlighted by saliency analysis and protein contextualization cannot be confirmed to be causative without further *in silico* and *in vitro* corroboration, although further validation in independent data will support a causal role. Second, traditional laboratory-based susceptibility testing can have high variance, especially for second-line drugs, introducing a potential source of error. Third, there is insufficient phenotypic data for certain anti-TB drugs (e.g. the novel agents bedaquiline and pretomanid, and second-line agents like ethionamide). Fourth, the non-causal mutation correlations observed in the MD-CNN boosted performance, but both the training and test data were enriched for multi-drug resistance. Further assessment of generalizability to a clinical setting with a low background prevalence of multi-drug-resistant *M. tuberculosis* is needed. Finally, additional computational resources would allow the inclusion of more loci of interest, likely augmenting the performance of the MD-CNN and SD-CNNs.

We believe this to be the first study to demonstrate the feasibility of interpretable, convolutional neural networks for prediction of antibiotic resistance in *M. tuberculosis*. Greater interpretability, reliability and accuracy make this model more clinically applicable than existing benchmarks and other deep learning approaches. Saliency mapping and protein contextualization also offer the possibility of creating hypotheses on mechanisms of anti-TB drug resistance to focus further research. Along with increasingly accessible WGS-capable infrastructure globally, machine-learning-based diagnostics may support faster initialization of appropriate treatment for MDR-TB, reducing morbidity and mortality, and improving health economic endpoints(*1*, *29*).

## Acknowledgements

We thank members of the Farhat lab for discussion and input. We are grateful to Dr. Peter Koo, Dr. Avika Dixit, and Greg Raskind for discussions regarding importance score calculation, validation analyses on the MD-WDNN, and CNN codebase proofreading, respectively. Computational resources and support were provided by the Orchestra High Performance Compute Cluster at Harvard Medical School, which is funded by the NIH (NCRR 1S10RR028832-01). AGG was supported by a National Institutes of Health NLM Training Grant T15LM007092 and NIH/NIAID F32AI161793. CHY was supported by the US-UK Fulbright Commission (USA/UK), the BUNAC Educational Scholarship Trust (UK), the Gavin and Ann Kellaway Research Fellowship (Auckland Medical Research Foundation, New Zealand), and the Royal Australasian College of Physicians Rowden White Fellowship (Australasia). MIG was supported by the German Research Foundation (GR5643/1-1). MF is supported by NIH/NIAID R01AI155765.

## Code Availability

Implementation of all models and data analysis can be found at: https://github.com/aggreen/MTB-CNN

## Materials and methods

### Sequence data

The training, cross-validation, and test datasets consist of a combined 23,049 *M. tuberculosis* isolates for which whole genome sequence data and antibiotic resistance phenotype data are available. The sequencing data are obtained through the National Center for Biotechnology Information database, PATRIC, and published literature,: 10,201 strains are in the “train” dataset (for training and cross-validation) (*6, 30–42*), 7,537 are in the holdout “test_1” dataset (for hold-out testing) (*31*, *43*–*47*), and the remaining 5,312 are in the holdout “test_GenTB” dataset (for hold-out testing) (*31*, *43*–*47*).

We process sequences in the train and test_1 datasets using a previously validated pipeline as described by Ezewudo et al. (2018), with modifications as elaborated by Freschi et al. (2020)(*42*, *48*). Briefly, reads are trimmed and filtered using PRINSEQ(*49*), contaminated isolates are removed using Kraken(*50*), and aligned to the reference genome H37Rv using BWA-MEM(*22*, *51*). Duplicate reads are removed using Picard, and we dropped isolates with less than 95% coverage of the reference genome at 10x coverage.

For the “test_GenTB” dataset, we prepare the sequencing data in accordance with the protocol in Groschel *et al*.(*52*), a different variant of the Ezewudo et al. pipeline.

With regard to curated genetic variants, the predictor sets of features for the multi-drug wide and deep neural network (MD-WDNN, see *Machine learning models* below) are processed as described by Chen et al. (2019)(*11*). Conversely, for the single-drug and multidrug convolutional neural networks (SD-CNN and MD-CNN, see *Machine learning models* below), only the FASTA files for the loci of interest are necessary.

### Antimicrobial resistance phenotype data

Culture-based antimicrobial drug susceptibility to 2-to-13 anti-TB drugs are available for all 23,049 isolates in the combined training, cross-validation, and test dataset, collated with quality control criteria described by Farhat et al. (2016)(*3*). Phenotypes (drug susceptibility test results) for isolates in the training and cross-validation dataset are from the ReSeqTB data portal, the PATRIC database, and manual curation of phenotypic data available in the literature(*6*, *30*–*42*). Phenotypes for the test dataset isolates are from data available in the literature(*31*, *43*–*47*). Each isolate’s phenotype is classified as resistant, susceptible, or unavailable, with respect to a combination of 13 possible first-line (rifampicin, isoniazid, pyrazinamide, ethambutol) and second-line drugs (streptomycin, ciprofloxacin, levofloxacin, moxifloxacin, ofloxacin, capreomycin, amikacin, kanamycin, ethionamide). (**Table 1**). In the hold-out test dataset, ethionamide and ciprofloxacin were excluded due to data missingness (0/2 resistant to ciprofloxacin; 12/25 resistant to ethionamide).

### Selecting input loci

The loci of the isolate sequences are selected from genes known or suspected to cause resistance based on previous models and experiments (**Table 1**). In order to incorporate regulatory sequences from the immediate genetic neighborhood, regions upstream from genes of interest are included. Loci were aligned to the H37Rv reference genome for comparison of coordinates and genome annotations are based on H37Rv coordinates from Mycobrowser(*53*).

### Machine learning models

The Multi-Drug (multi-task) Wide-and-Deep Neural Network (MD-WDNN) is described by Chen et al. (2019), and involves three hidden layers (256 ReLU), dropout, and batch normalization(*11*)

The Multi-Drug Convolutional Neural Network (MD-CNN) comprises two convolution layers (with filter size 12 nucleotides in length), one max-pooling layer, two convolution layers, one max-pooling layer, followed by two fully-connected hidden layers each with 256 rectified linear units (ReLU) (Table 1). This architecture is selected based on its performance, as defined by area under the receiver operator characteristic curve (AUC), compared to other architectures with fewer convolutional layers and differential filter sizes **(Supplementary Figure 1)**. Neither random nor cartesian grid search of optimal hyperparameters is conducted.

The MD-CNN is trained for 250 epochs via stochastic gradient descent and the Adam optimizer (learning rate of e^−9^). We select an optimal number of epochs based on minimizing validation loss (**Supplementary Figure 5**). The training is performed simultaneously using the resistance phenotype for all 13 drugs, hence the 13 nodes in the final output layer (Table 1), the output of each node corresponding to the sigmoid probability of the strain being resistant to the respective drug.

The MD-CNN’s loss function is adapted from the masked, class-weighted binary crossentropy function described by Chen et al. (2019)(*11*). This function addresses the dataset imbalance (missing resistance phenotypes for a varying number of drugs in any given isolate) by upweighting the sparser of the susceptible and resistant classes for each drug, and masking outputs where resistance status was completely missing.

The Single-Drug Convolutional Neural Networks (SD-CNNs) are thirteen individually trained convolutional neural networks, each trained to predict for only one drug, hence the output layer having size 1 instead of 13. Each SD-CNN is given only the input loci relevant to its particular antibiotic, resulting in different input sizes depending on the longest locus for each drug. The architecture for the SD-CNNs is otherwise identical to that of the MD-CNN. The SD-CNNs are initially trained for 150 epochs using stochastic gradient descent and the Adam optimizer (learning rate of e^−9^) and an optimal number of epochs for each SD-CNN is selected to minimize the validation loss **(Supplementary Table 4)**.

### Logistic regression benchmark

We build a logistic regression benchmark to evaluate the performance of our neural network models. For each of the 18 input loci used in the MD-CNN and SD-CNNs, we select all sites with a minor allele frequency of at least 0.1%, resulting in 3,011 sites across 23,049 genomes. Sites are then encoded using a major/minor allele encoding.

Using the same train/test partitioning as for the neural network models, we use GridSearchCV in Scikit-learn v.0.23.2(*54*) to select the optimal L2 penalty weight for a LogisticRegression classifier with balanced class weights. Hyperparameter search is performed for each drug independently, testing the values *C*=[0.0001, 0.001, 0.01, 0.1, 1]. After selecting the optimal L2 weight, we use five-fold cross-validation on the training set to assess the AUC, specificity, and sensitivity, selecting a model threshold that maximized the sum of specificity and sensitivity.

### Training and model evaluation

Five-fold cross-validation is performed five times to obtain the performance metrics – area under the receiver operator characteristic curve (AUC), sensitivity, specificity, and probability threshold (to maximize the sum of sensitivity and specificity) – and the 95% confidence intervals of the AUC values between the models.

Model performance on the hold-out test sets is evaluated using the probability threshold selected during training.

### Computational details

The MD-CNN is developed and implemented using TensorFlow 2.3.0 in Python 3.7.9 with CUDA 10.1(*55*–*57*). Model training is performed on an NVIDIA GeForce GTX Titan X graphics processing unit (GPU).

### Analysis of mis-predicted isolates

For each SD-CNN model, we compute the genetic distance (number of different sites) between all isolates in the training and test sets. Only the loci included in each SD-CNN model are incorporated in the calculation.

### Importance Score calculation

Importance scores are calculated using DeepLIFT v. 0.6.12.0, using the recommended defaults for genomics: “rescale” rule applied to convolutional layers, and “reveal-cancel” rule applied to fully connected layers. We use the H37Rv reference genome, which is sensitive to all antibiotics, as the baseline(*22*).

Importance scores for each isolate sequence are calculated relative to the H37Rv baseline. For our analysis of positions influencing antibiotic resistance prediction, we take the maximum of the absolute value of the scores at each position across all resistant isolates.

### Lineage variant analysis

We define lineage variants as those found in the Coll *et al*. or Freschi *et al*. barcode of lineage-defining variants(*58*, *59*). We further annotate any position in our 18 loci as lineage associated if that position has an identical distribution of major/minor alleles to any position in the Freschi *et al* barcode, excluding the position *1,137,518* which defines lineage 7 (not present in our dataset).

