## Supplementary figures and tables for "A convolutional neural network highlights mutations relevant to antimicrobial resistance in *Mycobacterium tuberculosis*"

**Supplementary Table 1: Comparison of models by predictive performance.** Mean AUCs for first-line drugs and mean AUCs for second-line drugs are compared between logistic regression and the MD-CNN (LR v MD-CNN), logistic regression and the WDNN (LR v WDNN) and so on. Welch's two-sample t-test is used to determine the statistical significance of the difference in mean AUCs between models. Benjamini-Hochberg p-value correction is performed for multiple testing. Note that ciprofloxacin, ethionamide, and levofloxacin were excluded from the list of second-line drugs because the original WDNN was not trained on these models.

| Model comparison | Drug group | Difference in mean AUCs [95% CI] | p | Benjamini-Hochberg p |
|---|---|---|---|---|
| **LR v MD-CNN** | First-line drugs | -0.026 [-0.047, -0.0039] | 0.032 | 0.055 |
| **LR v WDNN** | First-line drugs | -0.037 [-0.058, -0.016] | 0.0022 | 0.0067 |
| **LR v SD-CNN** | First-line drugs | -0.015 [-0.037, 0.006] | 0.18 | 0.2 |
| **WDNN v MD-CNN** | First-line drugs | 0.011 [-0.0015, 0.024] | 0.10 | 0.15 |
| **WDNN v SD-CNN** | First-line drugs | 0.022 [0.0096, 0.034] | 0.0014 | 0.0055 |
| **MD-CNN v SD-CNN** | First-line drugs | 0.010 [-0.0034, 0.024] | 0.16 | 0.19 |
| **LR v MD-CNN** | Second-line drugs | -0.055 [-0.088, -0.022] | 0.0031 | 0.0075 |
| **LR v WDNN** | Second-line drugs | -0.061 [-0.095, -0.028] | 0.0011 | 0.0055 |
| **LR v SD-CNN** | Second-line drugs | -0.029 [-0.064, 0.0059] | 0.12 | 0.16 |
| **WDNN v MD-CNN** | Second-line drugs | 0.0066 [-0.0055, 0.019] | 0.3 | 0.3 |
| **WDNN v SD-CNN** | Second-line drugs | 0.032 [0.016, 0.049] | 0.00044 | 0.0052 |
| **MD-CNN v SD-CNN** | Second-line drugs | 0.026 [0.0091, 0.042] | 0.0045 | 0.0090 |

**Supplementary Figure 1: Comparison of model architectures for MD-CNN.** Mean and standard deviation of cross-validation AUC for three multi-drug convolutional neural network architectures are shown. We performed five-fold cross validation using our training data for three architectures: a 2x conv-conv-pool architecture with filter size 12, a 2x conv-pool architecture with filter size 12, and 2x conv-pool architecture with filter size 21. The 2x conv-conv-pool architecture was chosen for further study.

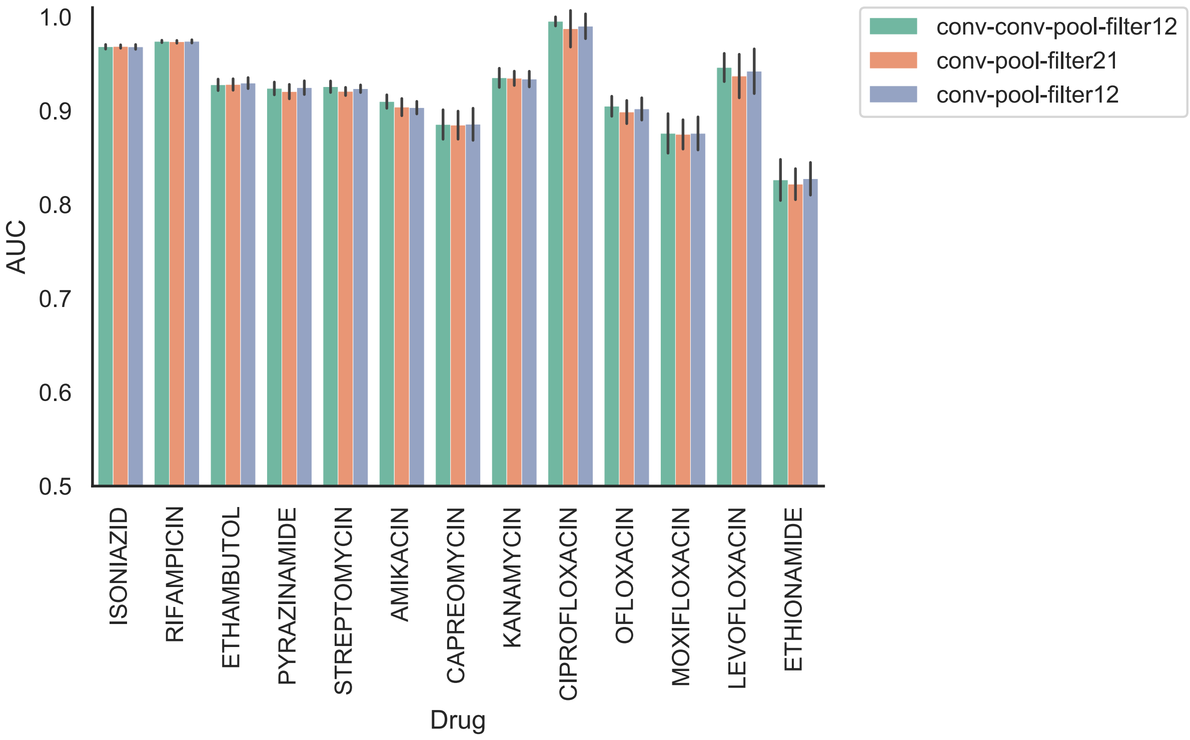

**Supplementary Figure 2:** schematic diagram of the single-drug convolutional neural network (SD-CNN). In the output layer, there is one node to compute a probability of resistance for the respective anti-TB drug. The input consisted of ‘10,201’ isolates (TB strains) for which there was resistance phenotype data for at least 2 anti-TB drugs; 5 for one-hot encoding of each nucleotide (4 dimensions, one for each nucleotide plus gaps); no. of nucleotides in the locus/loci of interest for the respective drug (selected as detailed in ‘Materials and methods’).

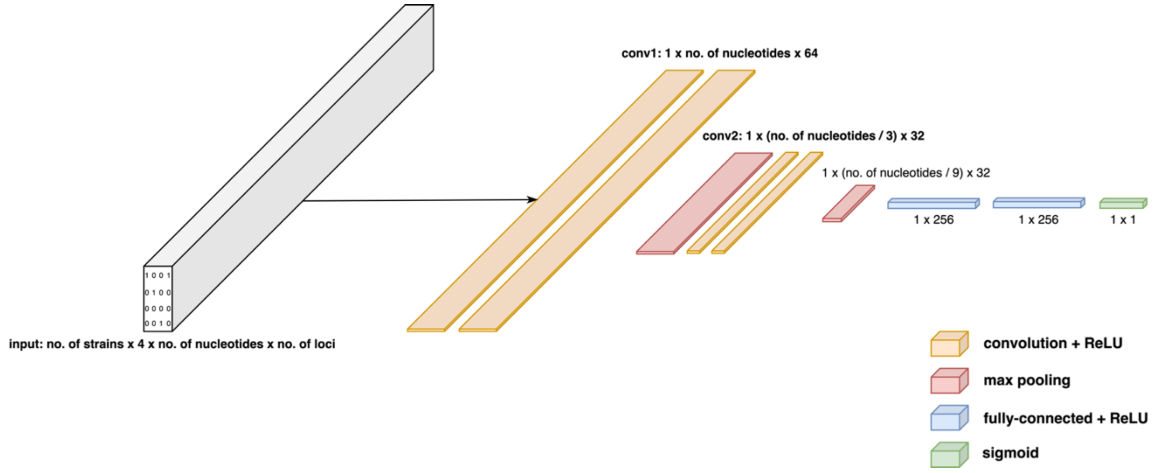

**Supplementary Figure 3: Comparison of false positives and negatives to training data phenotypes for SD-CNN.** Proportion of false negatives/positives that are nearest to resistant or sensitive isolates in the training dataset. For each falsely predicted test set isolate, we calculate the consensus phenotype (majority R or majority S) for the nearest isolate(s) in the training dataset. We differentiate between identical isolates (isolates with 0 mutational distance) and nearest non-identical isolates.

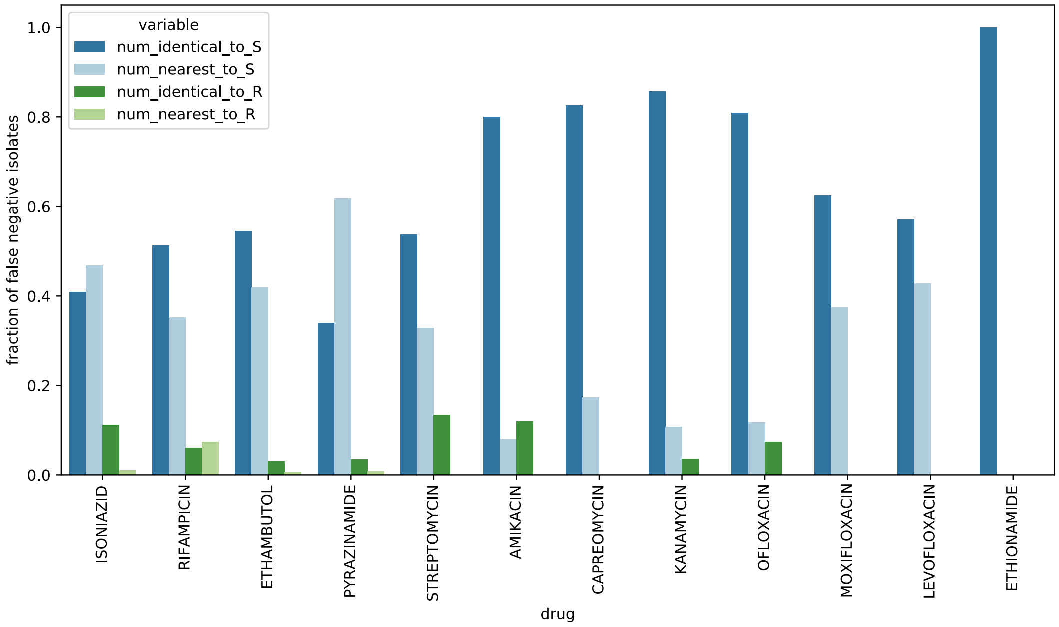

**Supplementary Table 2: High-importance sites according to DeepLIFT reflect a high proportion of known resistance-conferring variants.** Percent of high importance sites for each SD-CNN model that are classified as known to be involved in antibiotic resistance.

| *drug* | *percent known, top 1%* | *percent known, top 0.1%* |
|---|---|---|
| *AMIKACIN* | 6.84% | 63.64% |
| *CAPREOMYCIN* | 4.49% | 50.00% |
| *ETHAMBUTOL* | 5.37% | 50.00% |
| *ISONIAZID* | 9.35% | 46.15% |
| *KANAMYCIN* | 7.89% | 72.73% |
| *LEVOFLOXACIN* | 6.25% | 75.00% |
| *MOXIFLOXACIN* | 12.50% | 100.00% |
| *OFLOXACIN* | 8.51% | 100.00% |
| *PYRAZINAMIDE* | 62.26% | 80.00% |
| *RIFAMPICIN* | 55.26% | 85.71% |
| *STREPTOMYCIN* | 26.51% | 37.50% |

**Supplementary Table 3: Lineage-defining variants represent a small fraction of high importance positions in SD-CNN models.** The number of sites per SD-CNN model, the number of those sites that have been mutated (SNP or INDEL) at least once, the number of sites that are correlated with lineage-defining variants **(Methods)**, the number of hits (sites that comprise the top 0.1% of high-importance positions), and the number of hit sites that are correlated with lineage are shown.

| *drug* | *N sites* | *N sites mutated* | *N lineage sites* | *N hits* | *N lineage hits* | *% lineage sites* | *% lineage hits* |
|---|---|---|---|---|---|---|---|
| *AMIKACIN* | 12168 | 3997 | 3 | 121 | 2 | 0.08% | 1.65% |
| *CAPREOMYCIN* | 12168 | 3384 | 2 | 121 | 1 | 0.06% | 0.83% |
| *ETHAMBUTOL* | 20582 | 3468 | 19 | 205 | 11 | 0.55% | 5.37% |
| *ISONIAZID* | 14940 | 6161 | 15 | 149 | 5 | 0.24% | 3.36% |
| *KANAMYCIN* | 12168 | 3997 | 3 | 121 | 1 | 0.08% | 0.83% |
| *LEVOFLOXACIN* | 4859 | 1410 | 8 | 48 | 1 | 0.57% | 2.08% |
| *MOXIFLOXACIN* | 4859 | 1410 | 8 | 48 | 3 | 0.57% | 6.25% |
| *OFLOXACIN* | 4859 | 1410 | 8 | 48 | 4 | 0.57% | 8.33% |
| *PYRAZINAMIDE* | 17864 | 4820 | 10 | 178 | 4 | 0.21% | 2.25% |
| *RIFAMPICIN* | 7910 | 1955 | 9 | 79 | 4 | 0.46% | 5.06% |
| *STREPTOMYCIN* | 18252 | 4258 | 3 | 182 | 3 | 0.07% | 1.65% |

**Supplementary Figure 4: Rifampicin importance score plot for SD-CNN model.** Plot of maximum DEEPLIFT importance score across the rpoB gene (note that both *rpoB* and *rpoC* were included as inputs, only *rpoB* is shown here for resolution). The five highest importance scores not previously known to cause resistance are highlighted in purple and visualized on the rpoB protein structure (PDB ID 5UH5) along with all higher-scoring known variants.

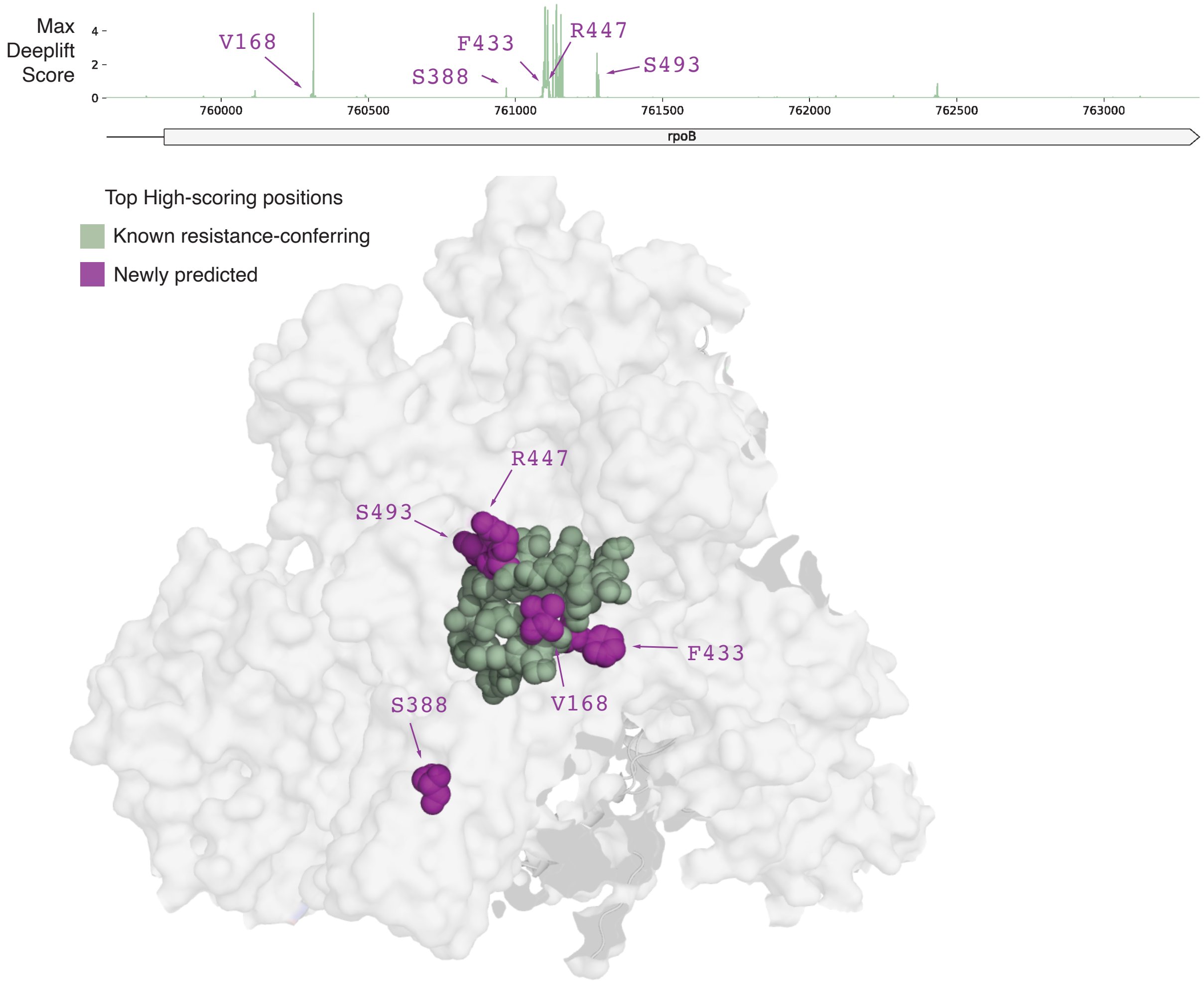

**Supplementary Figure 5: MD-CNN validation accuracy and loss during cross-validation.** Five-fold cross validation was performed on the MD-CNN to select the optimal number of epochs for trainings. The mean validation accuracy and validation loss was computer across the five splits. Final number of epochs selected was 150.

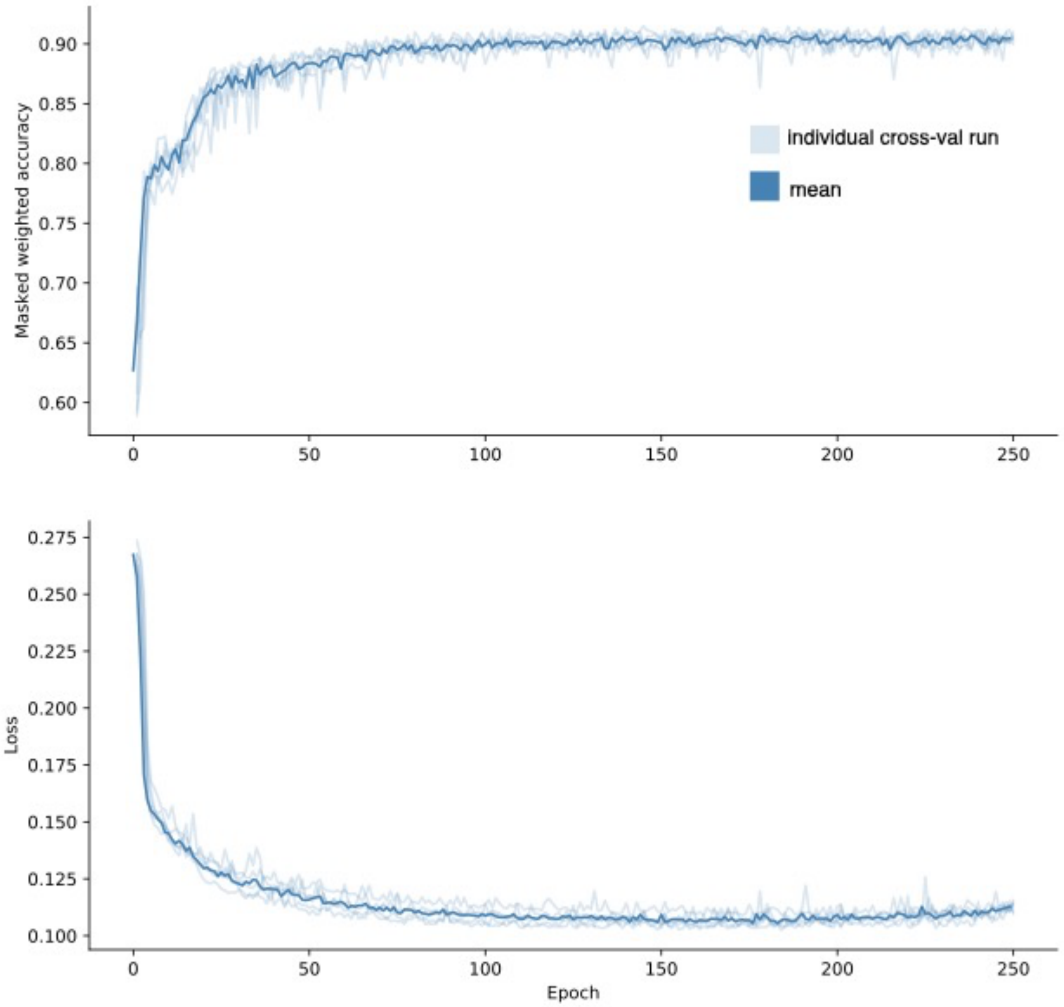
